# Cellular contractile forces are non-mechanosensitive

**DOI:** 10.1101/733303

**Authors:** Lea Feld, Lior Kellerman, Abhishek Mukherjee, Ariel Livne, Eran Bouchbinder, Haguy Wolfenson

## Abstract

Cells’ ability to apply contractile forces to their environment and to sense its mechanical properties (e.g. rigidity) are among their most fundamental features. Yet, the interrelations between contractility and mechanosensing, in particular whether contractile force generation depends on mechanosensing, are not understood. We use theory and extensive experiments to study the time evolution of cellular contractile forces and show that they are generated by time-dependent actomyosin contractile displacements that are independent of the environment’s rigidity. Consequently, contractile forces are non-mechanosensitive. We further show that the force-generating displacements are directly related to the evolution of the actomyosin network, most notably to the time-dependent concentration of F-actin. The emerging picture of force generation and mechanosensitivity offers a unified framework for understanding contractility.

**One Sentence Summary:** Cellular contractile forces are generated by rigidity-independent displacements that are determined by the time evolution of F-actin assembly.

## Main Text

The mechanical properties of the extracellular matrix (ECM) play critical roles in the most fundamental cellular processes (*1*). Matrix rigidity, for instance, affects cell polarization, migration, growth, death, and differentiation (*2–9*). Probing ECM mechanical properties (‘mechanosensing’) and feeding the outcome back into cells (‘mechanotransduction’) are processes that occur via integrin-based cell-matrix adhesions. These multi-molecular complexes bind the ECM at the extracellular side and attach to the actomyosin cytoskeleton at the intracellular side (see Fig. 1A). The cytoskeleton is composed of a network of actin filaments (F-actin) and myosin motors, where the latter actively generate contractile forces. It is well established that with increased ECM stiffness, larger contractile forces are transmitted to the matrix (*10–13*). The common view is that at any point in time, this transmission is regulated through the strength of the adhesions (*11*). The formation of large adhesions (classically referred to as ‘focal adhesions’) on stiff matrices is typically accompanied by the formation and growth of thick actin stress fibers that are attached to them (*4*). Collectively, these processes have dramatic effects on cellular properties and functions. Still, although actomyosin-based cellular contractility is regarded as a fundamental mechanosensing mechanism, a comprehensive understanding of both the contractile force generation process and mechanosensitivity, and their interrelations, is currently missing. In particular, whether the contractile force generation process is mechanosensitive, i.e. regulated through feedback from the cell’s probing/sensing of the rigidity, is not known.

**Fig. 1.**
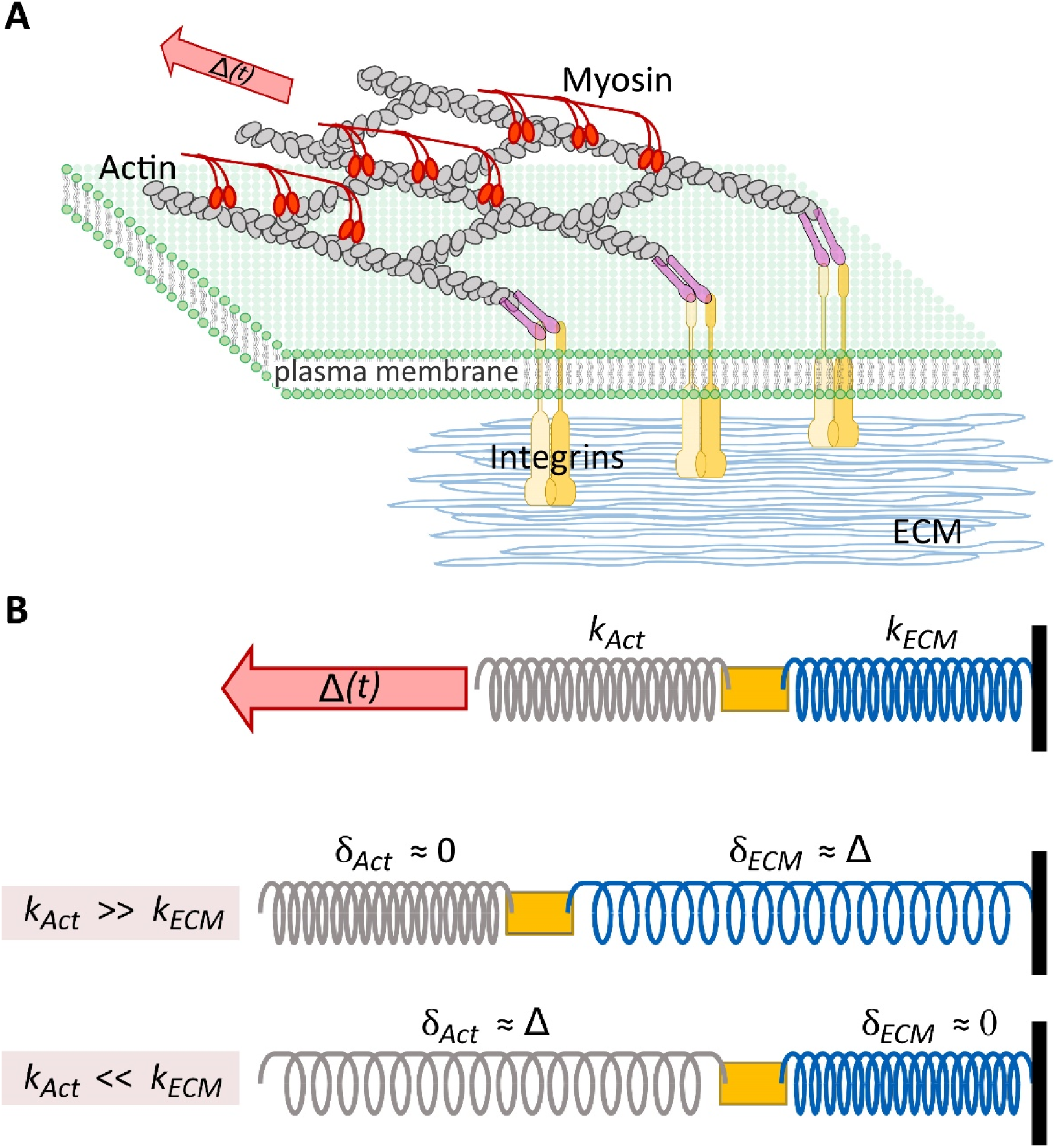
A simple model of cellular contractility. (A) A schematic of the cellular force generation machinery. that Shown are the integrin molecules, which are attached to the ECM at their extracellular side, and to actin structures (indirectly, through adapter proteins – purple rods) at the cellular side. Myosin motors that attach to the actin filaments generate a time-dependent displacement, Δ(*t*) (red arrow). (B) Top: An abstraction of panel A in which a spring of effective rigidity *k*_*ECM*_, representing the ECM, is connected in series to another spring of effective rigidity *k*_*Act*_, representing the actin structures. The contractile displacement Δ(*t*), which is applied to the actin structures, is also shown. Color code as in panel A. Middle: The degree to which the ECM or the actin structures are deformed depends on their relative rigidity. For *k*_*Act*_ ≫ *k*_*ECM*_, the myosin-generated contractile displacement Δ(*t*) is accommodated by ECM deformation, *δ*_*ECM*_ ≈ Δ, while the actin filaments composing the actin structures slide past each other as rigid objects with no internal deformation, *δ*_*Act*_ ≈ 0. Bottom: In the opposite limit (relevant for experiments performed on rigid ECM, e.g. plastic/glass plates), *k*_*Act*_ ≪ *k*_*ECM*_, the myosin-generated contractile displacement Δ(*t*) is accommodated by stretching the actin filaments composing the actin structures, *δ*_*Act*_ ≈ Δ, while the rigid ECM is not deformed, *δ*_*ECM*_ ≈ 0.

Here, we address the basic nature and time evolution of cellular contractile forces through a simple mathematical model combined with extensive experiments using cells adhering to micro-pillar arrays (*14*). We show that for a broad range of physiologically relevant ECM rigidities, the contractile force is proportional to the ECM rigidity as a result of simple mechanics. The proportionality factor is an intrinsic, cell-specific time-dependent contractile displacement that is *rigidity-independent*. Namely, we demonstrate that cellular contractile forces are themselves generically *non-mechanosensitive*. This fundamental observation implies that information about ECM rigidity is internally and simply encoded at each point in time in the magnitude of the contractile force. We further show that the time-dependent contractile displacement is determined by the time evolution of the F-actin concentration, indicating that the dynamics of myosin motors are not the rate-limiting factor in the cellular force generation process. The emerging picture of cellular contractility is shown to explain and unify various existing observations.

### A simple model of cellular contractility

The major component of the cellular contractile force machinery is actomyosin networks, which are made up of force transmitting actin structures and force generating myosin motors (Fig. 1A). For adherent cells, the generated contractile force is transferred across the plasma membrane to the ECM through integrins (Fig. 1A). To model how contractile forces are generated, we first assume that the myosin motors intrinsically generate a *time-dependent contractile displacement*, Δ(*t, k*_*ECM*_), which they impose at time *t* on the actin structures to which they attach and which may depend on the effective rigidity of the ECM, *k*_*ECM*_. Whether or not the contractile displacement depends on *k*_*ECM*_ is a major question to be addressed below. Note that the existence of a contractile displacement is in line with the fundamental property of myosin motors, which display a typical working stroke size (namely, distance) (*15*). Δ(*t, k*_*ECM*_) is accommodated by the displacement *δ*_*Act*_(*t*) of the actin structure and by the deformation *δ*_*ECM*_(*t*) of the ECM, i.e. Δ(*t*) = *δ*_*Act*_(*t*) + *δ*_*ECM*_(*t*). At any time *t*, the integrin adhesions are strong enough to withstand the forces applied to them.

The contractile force *F*(*t, k*_*ECM*_) satisfies the coarse-grained force balance equation *F*(*t, k*_*ECM*_) = *k*_*Act*_ *δ*_*Act*_(*t*) = *k*_*ECM*_ *δ*_*ECM*_(*t*), where *k*_*Act*_ is the effective rigidity of the actin structures (Fig. 1B, top panel; see also Supplementary Note 1 & Schwarz et al. (*16*)). This, in turn, implies that 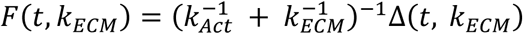 (for simplicity reasons we treat *k*_*Act*_ as a constant, although it is in fact time-dependent; see also Supplementary Note 2). In order to proceed, we further assume that for a broad range of physiologically relevant conditions (namely, relatively soft environments), the effective rigidity of actin structures is significantly larger than the rigidity of the ECM, i.e. *k*_*Act*_ ≫ *k*_*ECM*_. With this assumption, to be extensively tested below, we obtain

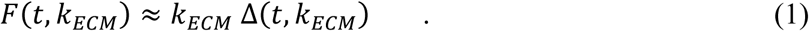

This equation shows that the dependence of the contractile force on the external rigidity may have two qualitatively different origins. First, the possible mechanosensitive contribution – if it exists – is contained within the contractile displacement Δ(*t, k*_*ECM*_). Second, and independently of the possible *k*_*ECM*_-dependence of Δ(*t, k*_*ECM*_), there exists also a non-mechanosensitive contribution – the overall pre-factor *k*_*ECM*_, which emerges from a linear force-displacement relation of the ECM. Consequently, Eq. (1) demonstrates that the mere existence of a dependence of *F*(*t, k*_*ECM*_) on the external rigidity *k*_*ECM*_ does not immediately indicate that the contractile force is mechanosensitive. In fact. Eq. (1) offers a clean procedure to extract the possible mechanosensitive part of the contractile force by plotting the force *F*(*t, k*_*ECM*_) normalized by *k*_*ECM*_, *F*(*t, k*_*ECM*_*)/k*_*ECM*_, a procedure that is followed in our experiments described below.

Note that *F*(*t, k*_*ECM*_*)/k*_*ECM*_ simply equals *δ*_*ECM*_(*t*), which implies that in the regime *k*_*Act*_ ≫ *k*_*ECM*_, the model predicts also *δ*_*ECM*_(*t*) ≈ Δ(*t, k*_*ECM*_) and *δ*_*Act*_(*t*) ≈ 0 (see Fig. 1B, middle panel). That is, in this case, the actin filaments/structures are predicted to experience no internal deformation, but rather to slide one relative to another as rigid objects (as they polymerize/de-polymerize at their edges), and the ECM deformation *δ*_*ECM*_(*t*) allows direct access to the myosin-generated collective contractile displacement Δ(*t, k*_*ECM*_). Finally, note that in the opposite regime, *k*_*Act*_ ≪ *k*_*ECM*_, relevant to cells adhering to glass/plastic plates, the model predicts *δ*_*Act*_(*t*) ≈ Δ(*t, k*_*ECM*_) and *δ*_*ECM*_(*t*) ≈ 0 (see Fig. 1B, bottom panel and Supplementary Note 2).

### Direct experimental tests demonstrate that cellular contractile forces are non-mechanosensitive

To test whether Δ(*t, k*_*ECM*_) in Eq. (1) depends on *k*_*ECM*_, i.e. whether it is mechanosensitive, we performed extensive experiments on cells adhering to arrays of fibronectin-coated flexible PDMS micro-pillars of 2μm diameter (see Methods and Fig. 2A). We monitored the cells from the very initial stages of attachment to the pillars, and as they were spreading, we tracked the cell-driven pillar deflections, which is simply *δ*_*ECM*_(*t*) (Fig. 2A). We used pillars with rigidities that vary by more than 15-fold, *k*_*ECM*_ = 2, 6, 31 pN/nm, equivalent to effective elastic moduli in the range *E*_*eff*_ ≈ 1.5 − 22 kPa (see Methods), thus allowing us to test the relevance of the relation *k*_*Act*_ ≫ *k*_*ECM*_ to a broad range of physiological conditions (*17*). The validity of this relation will not be assessed directly as there are significant uncertainties about the typical value of *k*_*Act*_; instead, we will indirectly assess it through the relation *δ*_*ECM*_(*t*) ≈ *F*(*t, k*_*ECM*_*)/k*_*ECM*_.

**Fig. 2.**
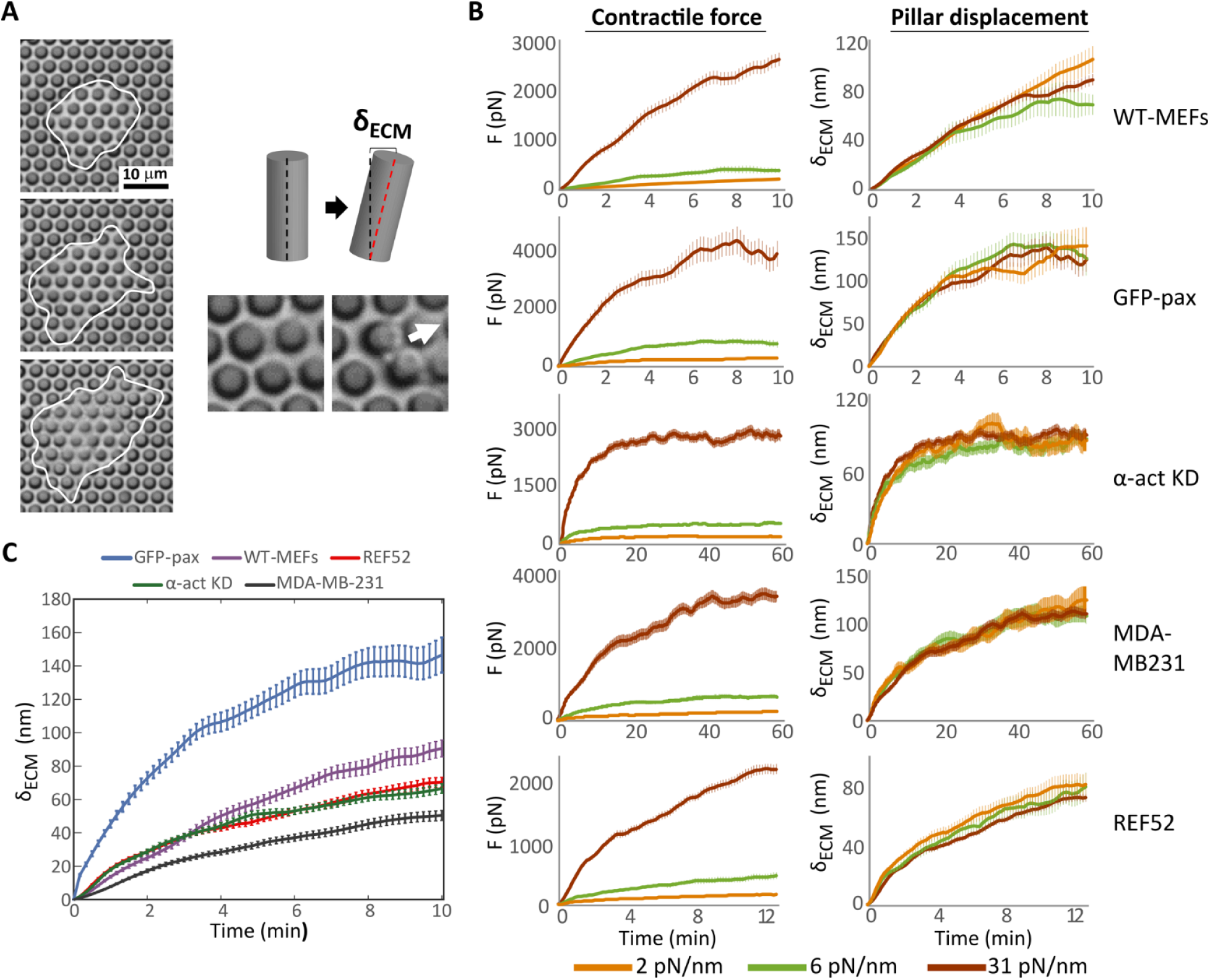
Pillar array experiments reveal a cell-specific, time-dependent, rigidity-independent contractile displacements. We used low-illumination brightfield time-lapse imaging to track live cells for several hours as they spread on the pillars to measure *δ*_*ECM*_(*t*). (A) Left: Frames from a time-lapse movie of a mouse fibroblast spreading on an array of 2μm diameter pillars coated with fibronectin (cell borders marked by the white line). Right: When the cell forms adhesions on the pillars, it pulls on them; pillar displacement (*δ*_*ECM*_) can be observed by tracking their tops over time. (B) Mean ± SEM of the time-dependent contractile forces (*F*(*t, k*_*ECM*_), left) and pillar displacement (*δ*_*ECM*_(*t*) = *F*(*t, k*_*ECM*_*)/k*_*ECM*_, right) on pillars of three widely different rigidities (corresponding to different pillar heights, the rigidity values appear on the legend). For all cell lines tested, *δ*_*ECM*_(*t*) = *F*(*t, k*_*ECM*_*)/k*_*ECM*_ collapsed onto a single master curve (right). n>50 pillars from >8 cells in each case. (C) Averages of the pillar displacement data from all three rigidities (panel B right) reveal cell-specific contractile displacements.

To test the generality of our model, we used five cell lines that displayed distinct phenotypic behaviors when adhering to soft and rigid matrices (Fig. S1): mouse embryonic fibroblasts (WT-MEFs), MEFs stably expressing paxillin-GFP (pax-GFP), MEFs with stable knockdown of α-actinin (α-act KD), rat embryonic fibroblasts (REF52), and the human breast adenocarcinoma cell line MDA-MB231. For all cell lines, we measured *δ*_*ECM*_(*t*) for multiple pillars from the initial rise of pillar displacement until the release (Fig. S2), averaged them, and superimposed the results for the widely different rigidities. The outcome, presented in Fig. 2B, is remarkable: for each cell line, the displacement function collapses on a master curve that is *independent* of the rigidity, despite the more than 15-fold variation in the latter. Thus, *δ*_*ECM*_ is time-dependent but *k*_*ECM*_-independent, and *F*(*t, k*_*ECM*_) ≈ *k*_*ECM*_ *δ*_*ECM*_(*t*) ≈ *k*_*ECM*_ Δ(*t*). This result, which indicates that the contractile force *F* trivially depends on *k*_*ECM*_ through a linear spring relation and hence is *non-mechanosensitive*, is one of the major results of this paper.

The contractile displacement, Δ(*t*), which is independent of *k*_*ECM*_, features generic properties that are cell line independent, but also quantitative differences between cell lines. In all cell lines, Δ(*t*) exhibits a gradual increase with time, until a plateau is reached on a time scale of 5-10 minutes. The exact time to reach the plateau and the plateau level itself do depend on the cell line (Fig. 2C). Moreover, while WT-MEFs and pax-GFP cells often released the pillars (i.e. decrease of the displacement/force) shortly after reaching the plateau, α-act KD and MDA-MB231 cells maintained the pillar displacements for much longer periods (tens of minutes on average) (Fig. 2B), and REF52 often showed a second increase in Δ(*t*) several minutes after reaching the plateau (Fig. S3).

### Non-mechanosensitive contractile displacements are directly related to the time-dependent concentration of F-actin

The most outstanding questions emerging from the findings presented above are related to the origin of the non-mechanosensitive, intrinsic time-dependent contractile displacement Δ(*t*), in particular, its typical time scale and plateau level. Since myosin motors drive contractile force generation, it is logical to assume that the displacement timescales and plateau levels are controlled by the gradual increase in myosin activity. However, myosin is constantly activated and recruited to actin filaments at the cell edge (*18*), and, moreover, myosin motors generate a quantum of contractile displacement (the working stroke size) on a time scale of milliseconds^15^, orders of magnitude faster than the time scale characterizing Δ(*t*). This may suggest that it is not the myosin motors per se, but rather the structural evolution of the actin network, to which the myosin motors attach, that might underlie Δ(*t*). Thus, we turned to characterize the evolution of the actin networks during force generation using the simplest quantity, i.e. the local (near pillar) concentration (number density) of F-actin, *C*_*F*−*actin*_(*t*).

To that end, we used three of the five original cell lines that expressed the F-actin reporter tdTomato-tractin (*19*) – WT MEFs, α-act KD cells, and MDA-MB-231 cells – and performed pillar displacement assays in parallel to tracking the fluorescence level of tractin around each pillar over time, which is a measure of *C*_*F*−*actin*_(*t*) (Fig. 3A). As the contractile displacement features a characteristic time of ∼ 10 min (cf. Fig. 2B), short time variations in *δ*_*ECM*_(*t*) and *C*_*F*−*actin*_(*t*) were smoothed out using a moving average window (see Methods); as shown in Fig. 3B, both curves appear highly correlated at all times. In fact, *δ*_*ECM*_(*t*) and *C*_*F*−*actin*_(*t*) of *individual pillars* are highly correlated across the different cell lines and rigidities, as shown in Fig. 3C. To test whether this relation persists to shorter time scales, we analyzed the *C*_*F*−*actin*_(*t*) on *δ*_*ECM*_(*t*) data without any smoothing (Fig. 3D). The results reveal that both *C*_*F*−*actin*_(*t*) and *δ*_*ECM*_(*t) synchronously oscillate* approximately every 4 minutes, regardless of cell type (Fig. 3E). In >80% of the cases when a rise in displacement was observed, it was accompanied by a simultaneous rise in *C*_*F*−*actin*_. Remarkably, the slightest increase in F-actin density was paralleled by an increase in the contractile displacement, and moreover, short-time scale disassembly of F-actin (decrease in tractin intensity) was paralleled by a drop in displacement (Fig. 3D, Fig. S4). These results were further substantiated by experiments in which *C*_*F*−*actin*_(*t*) has been significantly and promptly reduced using the actin polymerization inhibitor Latrunculin A, leading to the disappearance of contractile displacements, Δ(*t*) → *0* (Fig. S5).

**Fig. 3.**
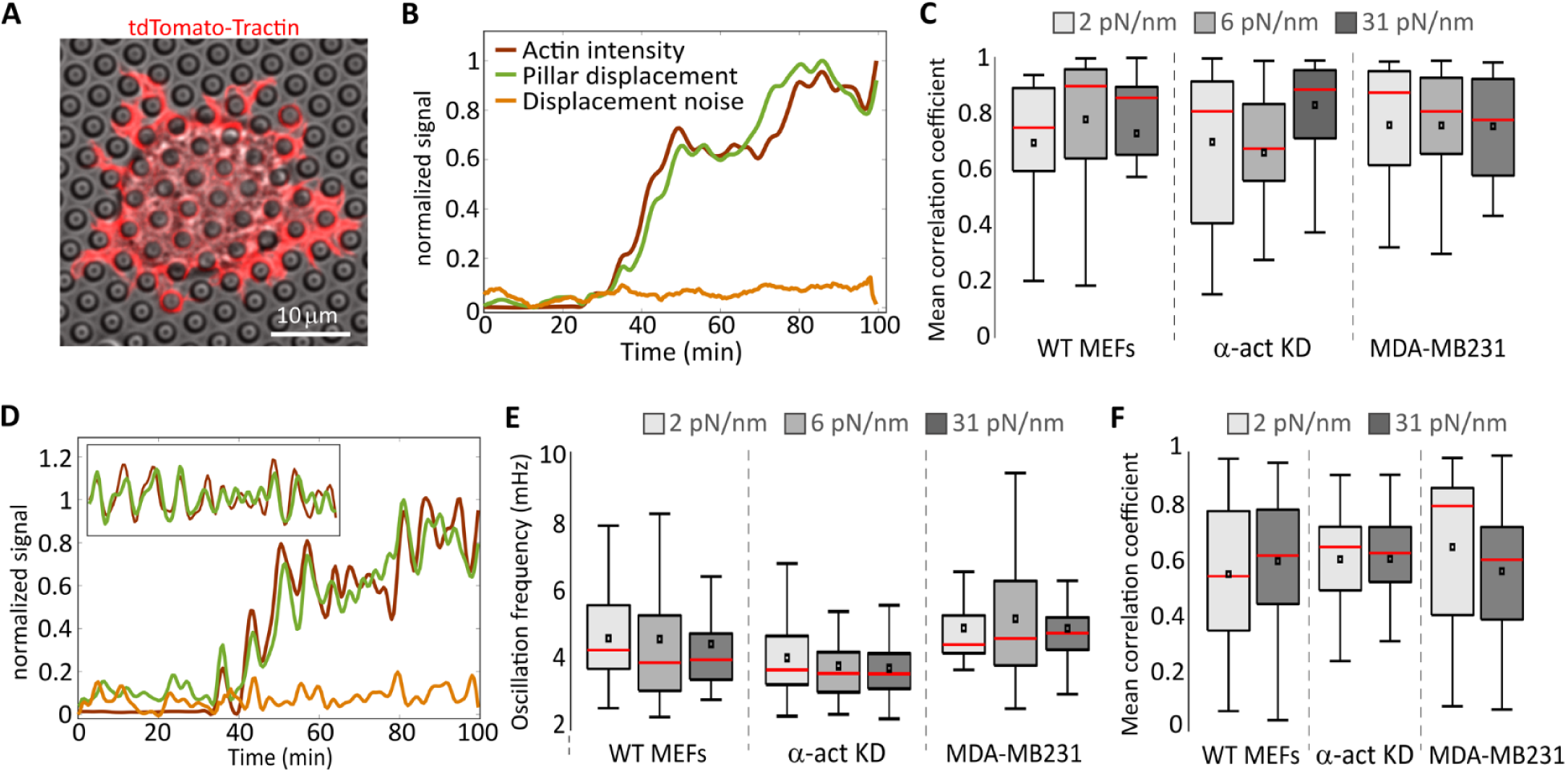
Correlation between F-actin density and pillar displacement. (A) Micrograph of a mouse fibroblast (WT MEF) expressing tdTomato-tractin and spreading on 2μm-diameter pillars. (B) Example of low-pass filtered curves of pillar displacement and of tractin intensity around the same pillar over time. (C) Mean correlation coefficients of pillar displacement and tractin intensity such as in panel B. n>30 from >5 cells in each case. The amplitude of the displacement noise, obtained by measuring the magnitude of the displacement (irrespective of its direction) of a pillar that was not in contact with the cell throughout the experiment, is added for reference. (D) Non-low-pass-filtered pillar displacement and tractin intensity over time curves reveal simultaneous oscillations in both. Inset shows the same data (starting from the initial rise of both signals) after subtraction of the low-pass filter curves in each case (i.e., minus the so-called DC component). Colors are as in panel B (see legend there). (E) Mean frequency of pillar displacement oscillations. The frequency was calculated using Fourier transform. Tractin oscillated at a similar frequency in all cases (not shown). (F) Mean correlation coefficients of actin and myosin density between the pillars.

Taken together, these observations support the simple yet quite remarkable relation:

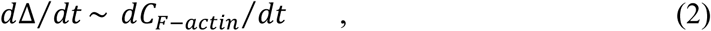

which suggests that the density of actin filaments essentially determines the contractile displacement generated by the actomyosin network. This implies, as hypothesized above, that myosin motors are not the rate-limiting factor in contractile force generation, i.e. that there are enough of them available at any point in time and that once attachment sites on the actin filaments are available, the myosin motors attach to them and generate displacement rapidly. This picture predicts that the densities of F-actin and myosin motors are highly correlated at any point in time, independently of ECM rigidity.

To test this prediction, we seeded cells on the pillar arrays with the two extreme rigidities (2 and 31 pN/nm) and fixed them after 30 min, during the contractile phase of spreading (*20*). We then stained the cells for F-actin and for phosphorylated (activated) myosin light chain (p-myosin). Using super-resolution microscopy (*21*), we observed large (micron-scale) actin structures between the pillars, which we classified as network-associated filaments, and small (nanometer-scale) unstructured filaments (Fig. S6A,B). p-Myosin was arranged in clusters, consistent with the appearance of myosin mini-filaments (*12, 22*) (Fig. S6C). We then tested the correlation between the density of the large F-actin structures and p-myosin between the pillars and found high positive correlation regardless of rigidity and cell type (correlation coefficients 0.6-0.7 in all cases; Fig. 3F), as predicted. Since each pillar in the collected images represents a different point in time during the displacement process, this indicates that whenever actin filaments are available, myosin motors operate on them.

### Cell-type dependence arises from structural differences in the actin cytoskeleton

The dependence of Δ(*t*) on *C*_*F*−*actin*_(*t*) implies that any possible cell-type dependence is encapsulated in the proportionality factor in Eq. (2). To test this, we measured the rate of change of *C*_*F*−*actin*_(*t*) and of *δ*_*ECM*_(*t*) (which measures Δ(*t*)) during the simultaneous short-time oscillations, and plotted the average *dδ*_*ECM*_*/dt* and *dC*_*F*−*actin*_*/dt* against each other to extract the proportionality factor. Note that by measuring the relative changes in F-actin concentration, we could disregard any differences in tractin transfection efficiency and in F-actin levels between cells. The resulting graphs indeed exhibit cell-type dependence; in particular, the proportionality factor of MDA-MB231 is significantly higher than that of the other two cell lines (Fig. 4A). This finding indicates that the degree to which the displacements follow changes in F-actin density varies between cell types.

**Fig. 4.**
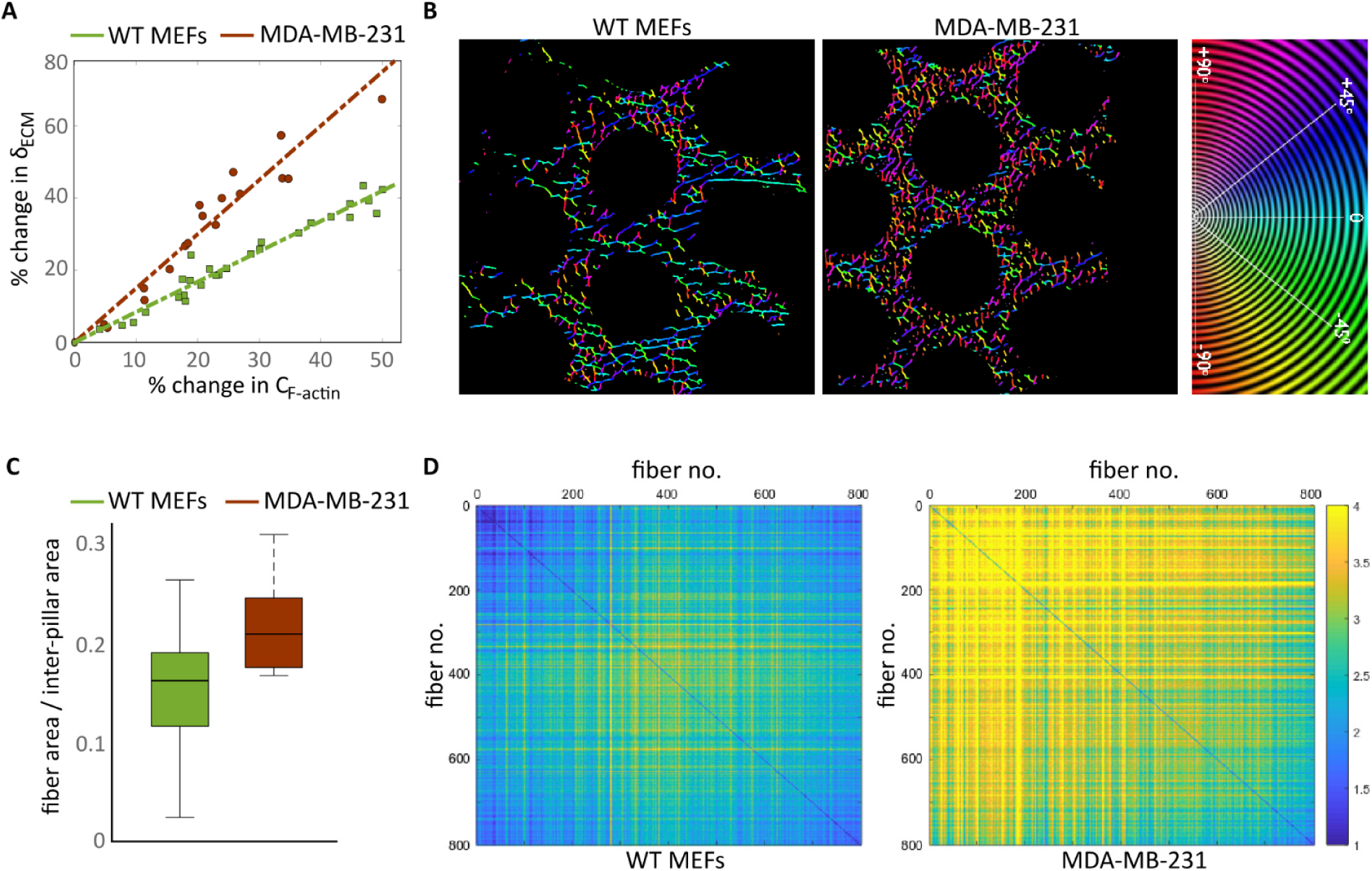
Structural differences in F-actin organization correlate with displacement response to *C*_*F*−*actin*_(*t*). (A) The rate of change (in percent) of *δ*_*ECM*_, which quantifies *d*Δ*/dt*, as a function of rate of change (in percent) of *C*_*F*−*actin*_, which quantifies *dC*_*F*−*actin*_*/dt*. Measurements were taken during each rise in the simultaneous oscillations (Fig. 3D), averaged for each rigidity (n>60 data points from >15 pillars from >4 cells in each case), and all data points from all three rigidities are plotted here for WT MEFs and MDA-MB-231 cells. For visual clarity, the α-act KD data (which are closer to those of WT MEFs than to these of MDA-MB-231 cells) are not shown. (B) Processed super-resolution images of large actin filaments at the cell edge color-coded for angles (see Methods for details). Only part of the cell edge is shown in each case; the right side of each image is outside of the cell. α-act KD cells displayed similar fiber distribution to that of WT MEFs (not shown). (C) Ratio between the area occupied by the large actin fibers and the inter-pillar area at the cell edge. MDA-MB-231 networks were ∼50% denser compared to WT MEFs (p-value < 0.001). (D) WT MEFs display highly parallel fibers compared to MDA-MB-231. Images such as those shown in panel B were analyzed as follows: the largest 800 fibers in each image were arranged according to size in ascending order and the inter-fiber slope differences were calculated for each fiber against all other fibers in each image. These differences are represented here by color-coded plots. The images shown are the average of 40 images in each case. Lower values (blue hues) represent parallel fibers. α-act KD showed a similar distribution to that of WT MEFs (not shown).

To shed light on this cell-type dependence, we utilized the super-resolution images of F-actin and p-myosin to characterize the assembled networks in WT MEF, α-act KD, and MDA-MB-231 cells (Fig. S6). We found that across the three cell lines, actin at the cell edges between pillars assembled into networks that occupied similar areas (0.2-0.35 μm^2^ per network) and similar complexity of branching (Fig. S6D). Interestingly, the number of p-myosin clusters per network in MDA-MB-231 cells was 30-40% lower than the other two cells lines (Fig. S6E), possibly related to the relatively low displacement rate (on timescales of tens of minutes) of MDA-MB-231 cells (Fig. 2C). This, however, cannot explain the higher proportionality factor in Eq. (2) (which relates to timescales of seconds) observed for MDA-MB-231 cells compared to the other cell lines, cf. Fig. 4A. The cell-type dependence of the proportionality factor instead implies that the degree to which changes in *C*_*F*−*actin*_ are translated into contractile displacement depends upon the geometrical properties of the assembled actin network. Indeed, analysis of the spatial organization of F-actin in the cells (Fig. 4B) revealed that whereas MDA-MB-231 cells display highly dense networks, WT MEFs typically form long, parallel, and more highly spaced filaments (Fig. 4C,D) (α-act KD cells were similar to WT MEFs, not shown). Thus, although additional work is required to quantitatively relate the geometry of assembled networks to the proportionality factor, the observed structural differences are likely to be related to the different proportionality factor in Eq. (2) of WT MEFs and α-act KD cells (i.e. different effective force transmission in response to changes in *C*_*F*−*actin*_).

### Predictions for well-spread cells in steady-state and consistency with existing literature

The experiments described above were performed on adherent cells that had not yet reached their well-spread steady-state with mature actin stress fibers that span a sizable area of the cell (*23*). On the other hand, our coarse-grained model makes no explicit reference to the specific actin structures that mediate the contractile displacement. Consequently, we expect the model to remain valid also for well-spread cells in steady state. To test this prediction, we used REF52 cells, which were previously shown to generate large forces at steady-state (*13*), and allowed them to attach to the pillars for 4 hours before imaging. These measurements showed that as in the case of early spreading, *δ*_*ECM*_(*t*) generated by well-spread REF52 cells collapses on a master curve that is rigidity-independent (Fig. 5A), strongly supporting the generality of our results. The time-dependent displacement featured a similar time scale (∼ 10 min) in both the steady-state (Fig. 5A) and early spreading (Fig. 2B) regime, possibly suggesting that the contractile displacement Δ(*t*) in the two regimes are intrinsically related. To test this, we plot one against the other in Fig. 5B. The result reveals that the two are (predominantly) linearly related to one another, demonstrating that the generated time-dependent displacement in the two regimes is in fact the *very same* function multiplied by a constant (the slope of the linear relation). The slope indicates that the early spreading displacements are 6-7-fold smaller than *δ*_*ECM*_(*t*) of the cells at steady state. This may imply that at steady state, when the cell edge extends and forms a new adhesion on a previously non-displaced pillar, the actin filaments that emanate from the new adhesion connect to pre-assembled stress fibers. Indeed, the deflected pillars in these cells were connected to thick actin stress fibers (Fig. 5C, Fig. S7) that were not observed in the early stages of spreading (Fig. 5D) (see also Supplementary Note 2).

**Fig. 5.**
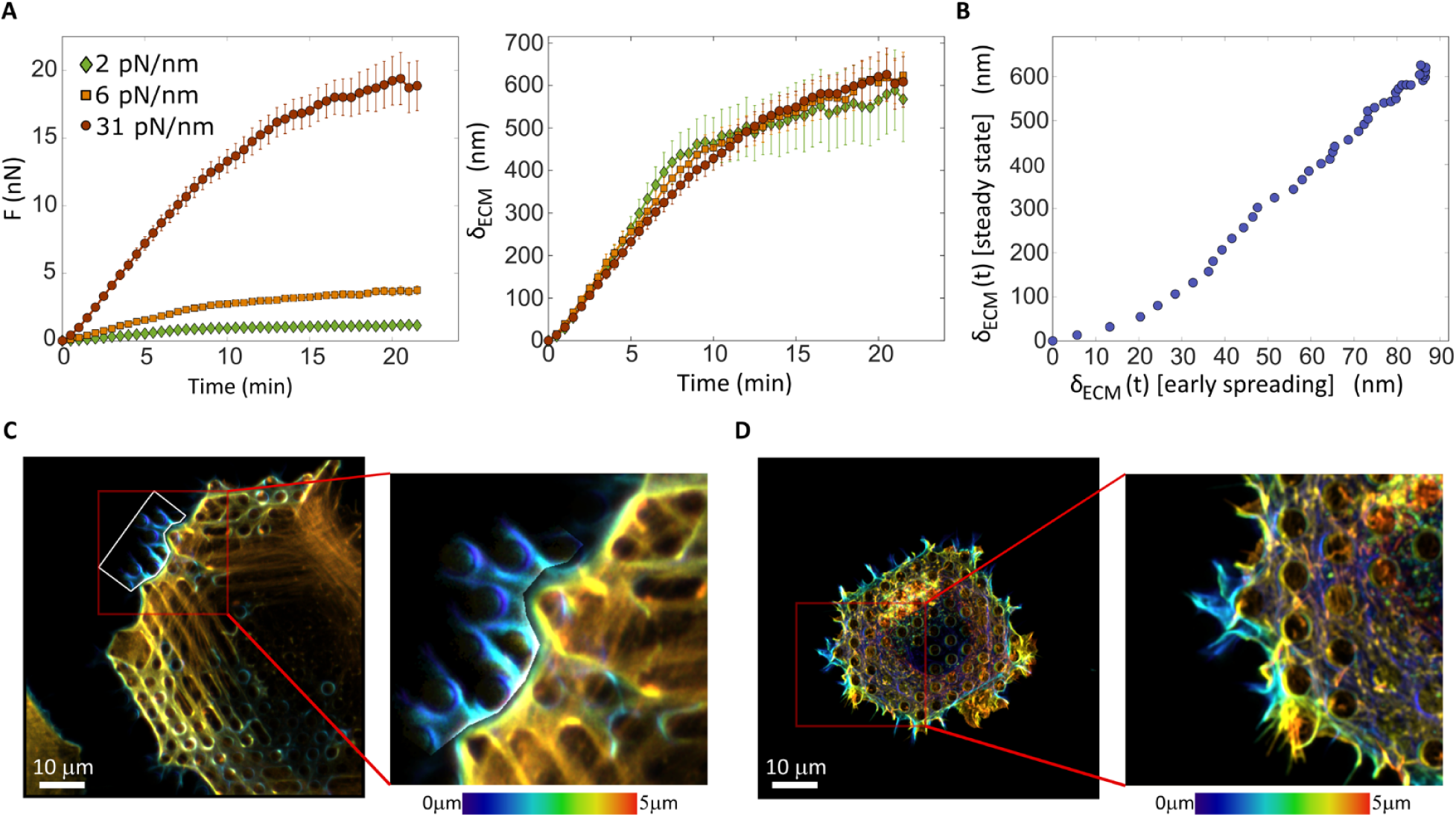
(A) *F*(*t, k*_*ECM*_) and *δ*_*ECM*_(*t*) = *F*(*t, k*_*ECM*_*)/k*_*ECM*_ curves (left and right, respectively) of REF52 cells under well-spread steady-state conditions. (B) *δ*_*ECM*_(*t*) of early spreading REF52 cells (cf. Fig. 2C) vs. *δ*_*ECM*_(*t*) of REF52 under well-spread steady-state conditions (panel B). (C-D) Z-stack projections of REF52 cells on 31pN/nm pillars stained for F-actin color-coded for depth; (C) Cell after 5h of spreading. The brightness of the region marked by white borders was enhanced for purpose of clarity. Thick actin stress fibers (yellow-orange hues) are observed 2-4µm above the fibers directly surrounding the pillar (blue hues). (D) Cell after 30min of spreading. No stress fibers are observed, and the actin structures are much less organized compared to steady-state condition (panel C).

This remarkable generality is also consistent with the previous results of Trichet et al.(*13*), who reported *F*(*t, k*_*ECM*_) for cells under well-spread steady state conditions over a broad range of pillars rigidities *k*_*ECM*_. To show this, we first plot 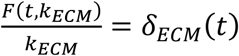 of Trichet et al.(*13*) in Fig. S8A,B, observing a collapse on a single master curve, as we predict and in agreement with our data in Fig. 5A. Moreover, in Fig. S8C we plot our *δ*_*ECM*_(*t*) (Fig. 5A) against that of Trichet et al.(*13*) (Fig. S8B), which once more reveals a linear relation, indicating that it is again the very same time-dependent contractile displacement (here, as both measurements were conducted under steady-state conditions, the slope is small, slightly below 2). Consequently, we conclude that cellular contractility under well-spread steady-state conditions is controlled by the very same intrinsic (non-mechanosensitive) displacement Δ(*t*), lending strong support to the generality of the emerging picture.

These results in the steady-state regime may have another intriguing implication. The independence of the intrinsic time-dependent contractile displacement Δ(*t*) on the rigidity *k*_*ECM*_ implies that Δ(*t*) is in fact also independent of the length of the stress fibers that are attached to the pillars through focal adhesions. This is immediately inferred from the clear dependence of the cell size, and consequently of the length of stress fibers, on the rigidity *k*_*ECM*_ as observed in Fig. 4 of Trichet et al. (*13*). On the other hand, one may assume that like in muscle cells (*24, 25*), myosin II motors apply their contractile displacement everywhere along the stress fiber, implying that Δ(*t*) increases with the length of the stress fiber. Our results (as well as those of Trichet et al. (*13*)) indicate that this is not the case, but rather that stress fibers of different lengths generate the *very same* contractile displacement Δ(*t*). This leads to the *activity localization hypothesis*, which suggests that myosin II motors apply their contractile displacement to a rather localized region of the stress fiber, and that the size of this region does not depend on the overall length of the stress fiber.

## Discussion

Our results reveal basic aspects of cellular force generation and mechanosensitivity, providing a novel framework to address various questions related to these important cellular processes. The starting point for our investigation is the widespread view that cellular contractile forces are mechanosensitive, i.e. that cells regulate these forces through sensing the rigidity of the ECM. This view is based on previous extensive observations that show that contractile forces depend on ECM rigidity: the larger the rigidity is, the larger cellular contractile forces are (*13, 26*–*28*). We show that cellular forces are generated through intrinsic non-mechanosensitive contractile displacements. That is, independently of the ECM rigidity, a particular cell type applies a well-defined time-dependent displacement to its environment (Fig. 2). This implies that the contractile force that is required to reach a certain displacement is proportional to the ECM rigidity, but that this dependence is purely mechanistic, i.e. resulting from a simple linear (Hookean) force-rigidity relation. Consequently, cellular contractile forces are themselves non-mechanosensitive.

These findings have major implications for our understanding of cellular force generation and mechanosensitivity. They show that while contractile cellular forces are not mechanosensitive, information about ECM rigidity is simply encoded in them. Hence, any mechanosensitive cellular process should be sensitive to the contractile force (e.g. its instantaneous magnitude or time rate of change), feeding back the rigidity information into other cellular processes. Moreover, as our findings show that the non-mechanosensitive force generation process emerges from intrinsic time-dependent displacements, any attempt to understand cellular force generation should focus on the intrinsic time-dependent displacements. As a first step towards achieving this important goal, we showed that the time-dependent contractile displacement is directly and universally related to the time evolution of actomyosin networks. In particular, it is shown to be causally related to the density of the actin filaments near the adhesion sites, and the degree to which changes in this density are translated into displacement is shown to depend on the cell-type specific spatial organization and structure of the networks formed by these filaments (Figs. 3-4).

Our results reveal a large degree of universality in the time-dependent contractile displacement. In particular, we show that very different cellular states – those pertaining to the early spreading process and those pertaining to steady-state spreading conditions – give rise to essentially the very same time-dependent contractile displacement, differing only by an overall amplitude (Fig. 5). These results are fully consistent with the experimental results of Trichet et al. (*13*), whose reanalysis also demonstrated non-mechanosensitive contractile displacements that differ by only a multiplicative constant from our own measurements in the well-spread steady-state (for the same cell type, here REF52). Our results also strongly support the relation *k*_*Act*_ ≫ *k*_*ECM*_ (used to derive Eq. (1)), which indicates that over a broad range of conditions, the actin structures involved in the force generation process are much stiffer than the ECM and do not deform at all during the process. This is consistent with previous *in vitro* measurements that showed that actin networks can be up to 40-fold stiffer than the loads that they bear (*29*).

The biophysical picture emerging from our findings is consistent with other available observations in the literature, and hence provide a unifying picture of cellular contractility. For example, while the inferred relation *k*_*Act*_ ≫ *k*_*ECM*_ is relevant to a broad range of physiological conditions, experiments performed on cells adhering to ultra-stiff substrates (e.g. plastic/glass) are in the opposite regime of *k*_*Act*_(*t, k*_*ECM*_) ≪ *k*_*ECM*_. Under these conditions, where the ECM cannot be deformed, we predict that the displacement Δ(*t*) is accommodated by stretching the actin structures, *δ*_*Act*_(*t*) ≈ Δ(*t*) > 0. Indeed, in laser ablation experiments applied to cells adhering to glass plates (*30*), stress fibers are observed to instantaneously retract by an amount comparable to the saturation level of *δ*_*ECM*_(*t* → *∞*) ≈ Δ(*t* → *∞*) observed in Fig. 5A and Fig. S8B (see also Supplementary Note 2). This observation further supports the major result that the contractile displacement Δ(*t*) is an intrinsic, non-mechanosensitive property of cells (independently of ECM rigidity, and independently of whether the displacement is accommodated by the ECM or by F-actin structures). It is important to stress that our use of ECM pillars whose effective rigidity varies with their height alone, keeping their material and surface properties entirely fixed, enables us to cleanly isolate the effect of the external rigidity, ruling out any possible intervening surface chemistry effects.

Previous studies of force regulation as a function of ECM rigidity have given rise to the ‘integrin clutch model’, which predicts that contractile forces depend on the number of attached clutches (integrins) to the moving, polymerization-driven, actin fibers (*31–33*). Our results show that the non-mechanosensitive contractility critically depends on the density of F-actin, as well as its spatial organization, making no explicit reference to the polymerization-driven flow of actin. As we have not measured the number of engaged clutches at each time-point during the contractile displacement process (nor is it possible to the best of our knowledge to accurately perform such measurements), the relation between the density of F-actin and adhesion dynamics should be further explored (e.g. in light of the common view that F-actin is recruited at the adhesion sites).

Our findings also give rise to various important questions. First, there is a need to go beyond the discovered *d*Δ(*t*)/*dt* ∼ *dC*_*F*−*actin*_(*t*)/*dt* relation, which in itself should be further studied, in order to understand the origin of some of the properties of Δ(*t*). In particular, the activity localization hypothesis associated with Δ(*t*) should be further tested in future work, together with the origin of the characteristic time scale and plateau levels of Δ(*t*), including their cell-type dependence. In addition, the mechanism behind the short-time oscillatory behavior of F-actin (characterized by a time of ∼ 4 min) should be clarified, potentially in relation to the cyclic activation/de-activation of actin polymerization factors, as was recently observed in contracting secretory vesicles (*34*). The relation between this oscillatory time scale and the lifetime of adhesions, during force loading (*35, 36*), including the turnover time of proteins within them, should be further explored. Finally, our main result, *F*(*t, k*_*ECM*_) ≈ *k*_*ECM*_ Δ(*t*), shows that mechanosensing is driven by cells through Δ(*t*) and that the sensing itself is done through the contractile force *F*(*t, k*_*ECM*_). How the latter feeds back into various internal cellular processes should be further explored with particular focus on the time-dependent accumulation of rigidity signals, rather than exclusively on their magnitude. This may shift our view of mechanosensitivity toward the dynamic nature of this process.

## Materials and Methods

### Cells culture and reagents

WT MEFs (RPTPα^+/+^ cells) (*37*) and α-act KD MEFs (*38*) were a kind gift from Mike Sheetz (MBI Singapore & The University of Texas Medical Branch); REF52 cells (*39*) and pax-GFP cells (Ilk^f/f^ fibroblasts stably expressing paxillin–EGFP) (*40*) were a kind gift from Benny Geiger (Weizmann Institute of Science); MDA-MB-231 cells (*41*) were a kind gift from Yuval Shaked (Technion). All cells were cultured at 37° C in a 5% CO2 incubator in Dulbecco’s Modified Eagle Medium (DMEM) supplemented with 10% fetal bovine serum, and 100 IU/ml Penicillin-Streptomycin (all reagents were from Biological Industries).

### Plasmids and transfections

The YFP-Tpm2.1 plasmid was a kind gift from Peter Gunning (The University of New South Wales). The tdTomato-tractin plasmid was a kind gift from Mike Sheetz (MBI Singapore & The University of Texas Medical Branch). Transfections were carried out 1 day before measurements using the NEPA21 Electroporator (Nepa Gene) according to the manufacturer’s instructions, with ∼ 10^6^ cells per reaction and 10 μg DNA.

### Pillar and soft gel fabrication

In the pillar experiments, the external rigidity *k*_*ECM*_ is controlled by varying the height of the pillars, while keeping their cross-sectional area and intrinsic chemical properties fixed. All pillars had a diameter of 2 µm, and heights of 5.3, 9.4, or 13.2 µm. We used 2 *μ*m diameter pillars as these can be used to measure the long-term time-dependent forces that are generated after initial formation and reinforcement of the adhesions(*10, 12*). The center-to-center spacing between pillars was 4 *μ*m. Pillar bending stiffness, *k*_*ECM*_, was calculated by Euler–Bernoulli beam theory:

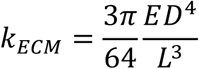

where D and L are the diameter and length of the pillar, respectively, and E is the Young’s modulus of the material (=2 MPa for the PDMS used here).

Using a common relation to estimate an effective elastic modulus that corresponds to a given rigidity 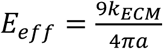, where *a* is the radius of the pillars (*42*), we obtain effective elastic moduli in the range *E*_*eff*_ ≈ 1.5 - 22 kPa. The latter is well within the relevant physiological regime, ranging from endothelial tissues to cartilage (*17*), thus allowing us to test the relevance of the relation *k*_*Act*_(*t, k*_*ECM*_) ≫ *k*_*ECM*_ to a broad range of physiological conditions.

Pillar fabrication was done by pouring PDMS (mixed at 10:1 with its curing agent, Sylgard 184; Dow Corning) into silicon molds (fabricated as previously described(*10*)) with holes at fixed depths and distances. The molds were then placed, face down, onto glass-bottom 35 mm dishes (#0 coverslip, Cellvis) which were incubated at 65°C for 12h to cure the PDMS. The molds were next peeled off while immersed in ethanol to prevent pillar collapse. The ethanol was replaced with PBS, and human plasma full-length fibronectin (Merck) was added to the dish at a final concentration of 10 µg/µl for a 1h incubation at 37°C. Next, residual fibronectin was washed away by replacing the buffer to HBSS buffer (Biological Industries) supplemented with 20 mM HEPES (pH 7.2).

1 and 40 kPa substrates (Supplementary Fig. 1) were fabricated by using Sylgard 52-276, at a ratio of 1:1.1 and 1:3.2, respectively, according to the measurements performed by Ou et al. (*43*).

### Pillar displacement measurements

One day prior to the pillar experiments, cells were sparsely plated to minimize cell-cell interactions prior to re-plating. The following day, cells were trypsinized, centrifuged with growth medium, and then resuspended and pre-incubated in HBSS/Hepes at 37°C for 30 min prior to the experiment. Cells were then spread on the fibronectin-coated pillars. In all cases, we made sure that the cells were isolated when plated on the substrates.

Time-lapse imaging of cells spreading on the pillars was performed using an inverted microscope (Leica DMIRE2) at 37°C using a 63x 1.4 NA oil immersion objective. Brightfield images were recorded every 10 seconds with a Retiga EXi Fast 1394 CCD camera (QImaging). The microscope and camera were controlled by Micromanager software (*44*). For each cell, a movie of 1-3 hours was recorded. To minimize photo-damage to the cells, a 600 nm longpass filter was inserted into the illumination path.

Tracking of pillar movements over time was performed with ImageJ (National Institutes of Health) using the Nanotracking plugin, as described previously(*12*). In short, the cross-correlation between the pillar image in every frame of the movie and an image of the same pillar from the first frame of the movie was calculated, and the relative x- and y-position of the pillar in every frame of the movie was obtained. To consider only movements of pillar from their zero-position, we only analyzed pillars that at the start of the movie were not in contact with the cell and that during the movie the cell edge reached to them. Drift correction was performed using data from pillars far from any cell in each movie. For each pillar, the displacement curve was generated by Matlab (MathWorks).

Parallel measurements of tdTomato-tractin intensity and pillar displacements were performed on a Zeiss LSM800 confocal microscope using a 63x 1.4NA objective at 37°C. Images were taken every 30 seconds and measurements of pillar movements were performed as described above. Measurements of tdTomato-tractin intensity were taken around each pillar of interest in a circular area with a radius of 2µm. Smoothing of the data in Fig. 3B was done using a moving average with a window size of 30 frames (5 minutes).

Analyses of correlation between changes in F-actin density and in pillar displacement during the short-term oscillations (Fig. 3D) were performed only on displacements that began below the noise level (∼ 20nm).

### Fluorescence microscopy

For immunofluorescence microscopy, cells were plated on fibronectin-coated pillars, fixed with 4% paraformaldehyde, and permeabilized with 0.1% Triton X-100. Immunolabeling was performed with primary antibodies against p-myosin (Abcam) overnight at 4°C, and with Alexa-488 or Alexa-555-conjugated secondary antibodies for 1 h at room temperature. Images were taken with a Zeiss LSM800 confocal microscope using a 63x 1.4NA objective.

For super-resolution analyses (Super-Resolution Radial Fluctuations; SRRF(*21*)), a region of interest (ROI) of 12µm × 12µm was imaged using the Airyscan function on the LSM800 confocal microscope. A series of 25 images was taken at 5fps; these were processed using the Airyscan algorithm using the ZEN blue software (Zeiss), followed by further processing using the SRRF algorithm(*21*). The trainable Weka segmentation plugin for FIJI (*45*) was then used to define the actin fibers between the pillars (Fig. 4), and thresholding was used to define the p-myosin clusters above background fluorescence levels. Fiber orientation analysis was done using the Ridge Detection and OrientationJ plugins for FIJI.

### Statistical analysis

Matlab (MathWorks) was used for data analysis and graph plotting. All ensemble average pillar displacement curves are shown with error bars representing the standard error of the mean (SEM). In the boxplots displayed in the figures the central mark indicates the median, the bottom and top edges of the box indicate the 25th and 75th percentiles, respectively, and the whiskers extend to the most extreme data points not considered outliers; outliers are not shown.

## Acknowledgments

We are grateful to Yuri Lubomirsky (Weizmann Institute of Science) for discussions related to Eq. (1), to Noam Kaplan (Technion – Israel Institute of Technology) for discussions on data analysis, and to Sasha Bershadsky (Mechanobiology Institute, Singapore) for discussions related to theoretical aspects of this work. H.W. acknowledges support from the Israel Science Foundation (1738/17) and from the Rappaport Institute. H.W. is an incumbent of the David and Inez Myers Career Advancement Chair in Life Sciences. E.B. acknowledges support from the William Z. and Eda Bess Novick Young Scientist Fund and from the Harold Perlman Family.

## Author contributions

L.F. performed the experiments and analyzed the data; L.K. and A.M. assisted in data analysis and pillar experiments, respectively; A.L. and E.B. conceived and developed the theory; E.B and H.W. conceived and planned the experiments; E.B. and H.W. wrote the manuscript. All authors discussed the results and contributed to the final manuscript.

