## Supplementary Information for "Cellular contractile forces are non-mechanosensitive"

### Cellular contractile forces emerge from time and F-actin dependent non-mechanosensitive displacements

### Supplementary Information

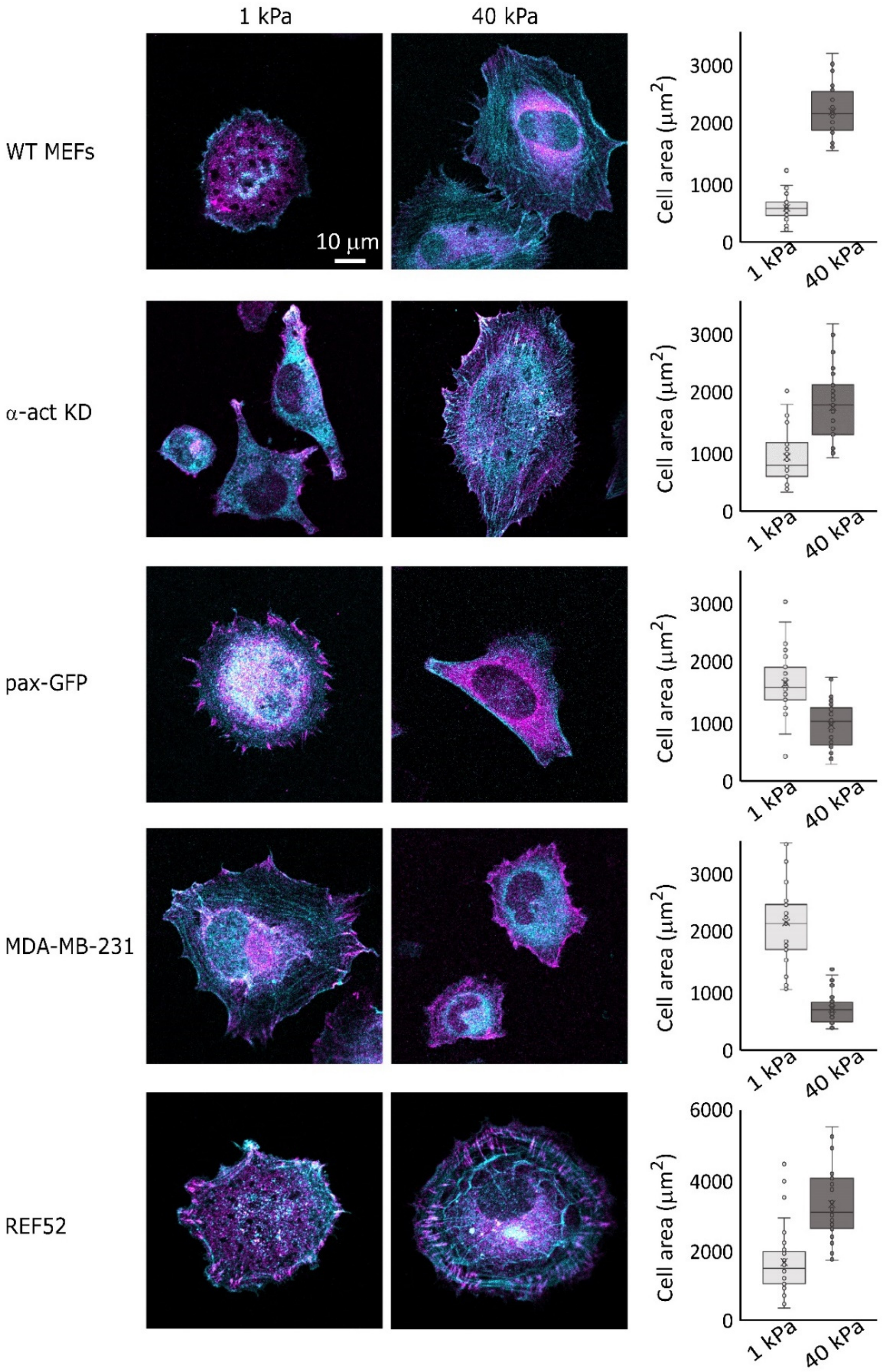


**Fig. S1.** Representative images of all 5 cell lines plated on 1 and 40 kPa continuous gels for 3h and stained for vinculin (magenta) and F-actin (cyan). Whereas WT-MEFs, α-act KD cells, and REF52 display larger adhesions and more organized stress fibers on stiff vs. soft, the opposite is observed for pax-GFP cells and MDA-MB-231 cells. The right boxplots show the differences in cell area on the different rigidities (n>30 cells in each case).

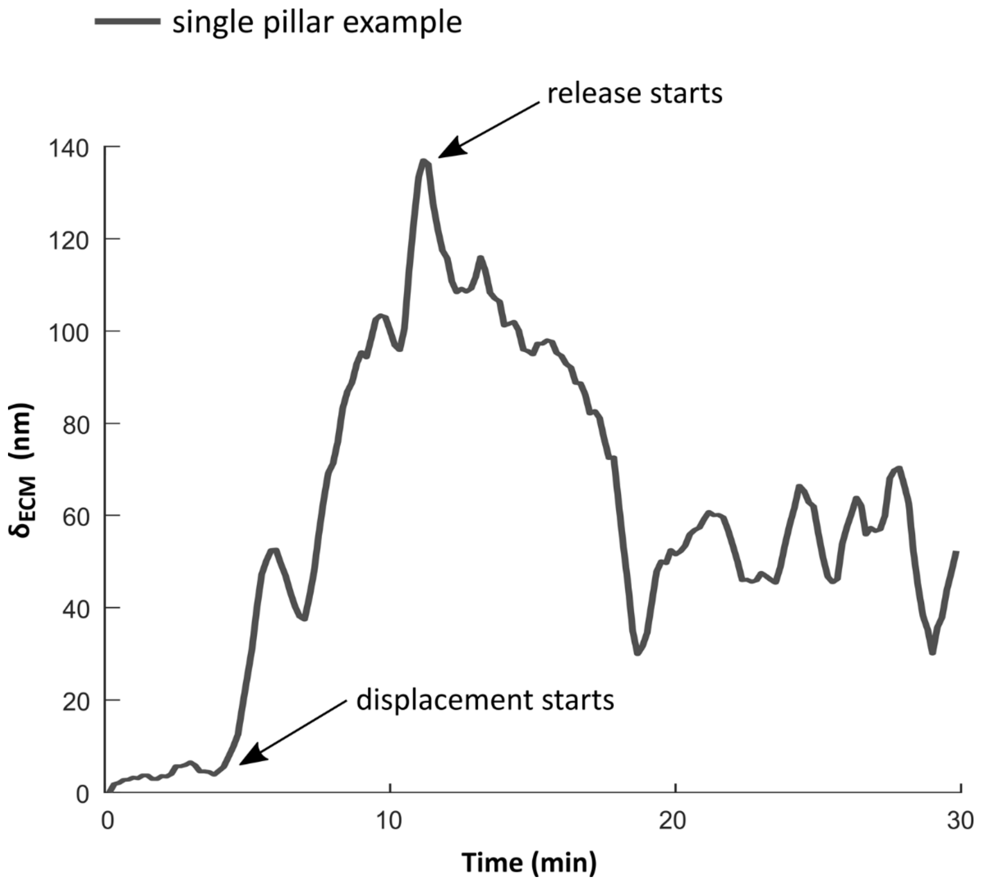


**Fig. S2.** Example of a single pillar displacement curve. The beginning of the release was defined by the time point of maximal displacement.

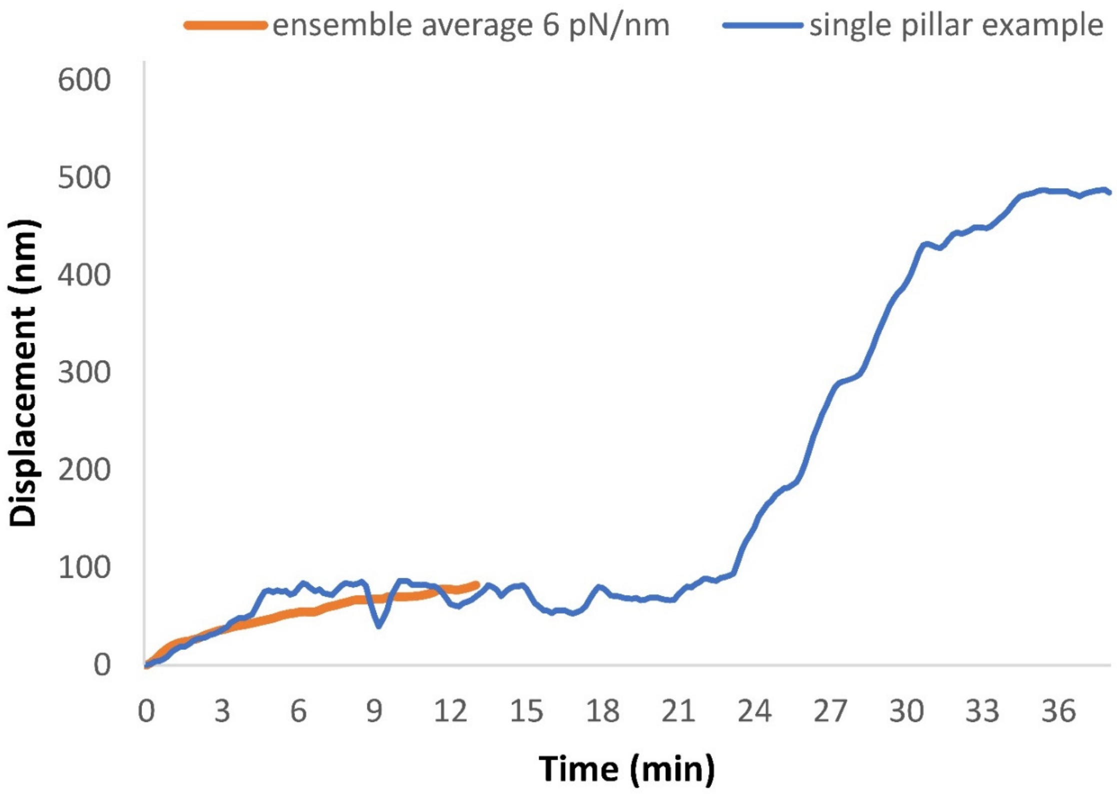


**Fig. S3.** Example of a single pillar displacement curve of a REF52 cell showing a second rise in displacement several minutes after an initial plateau was reached. The orange curve is the same as shown in Fig. 2B.

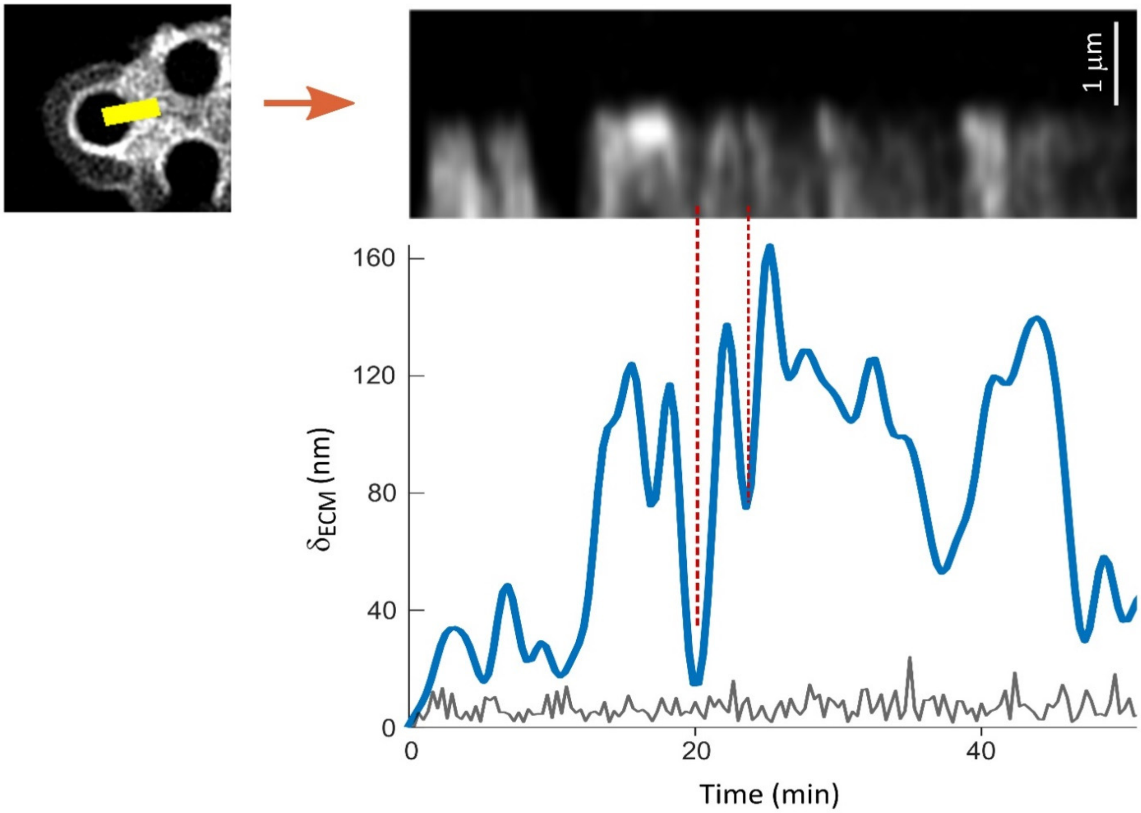


**Fig. S4.** Kymograph of tractin intensity taken from the edge of a single pillar (yellow rectangle in the left image was used to generate the kymograph) aligned with the displacement graph of the same pillar. Note the example of a single period between the dotted lines with simultaneous rise and drop in tractin intensity and in $\delta_{ECM}$. Grey curve is the displacement noise (pillar that was not in contact with the cell throughout the experiment).

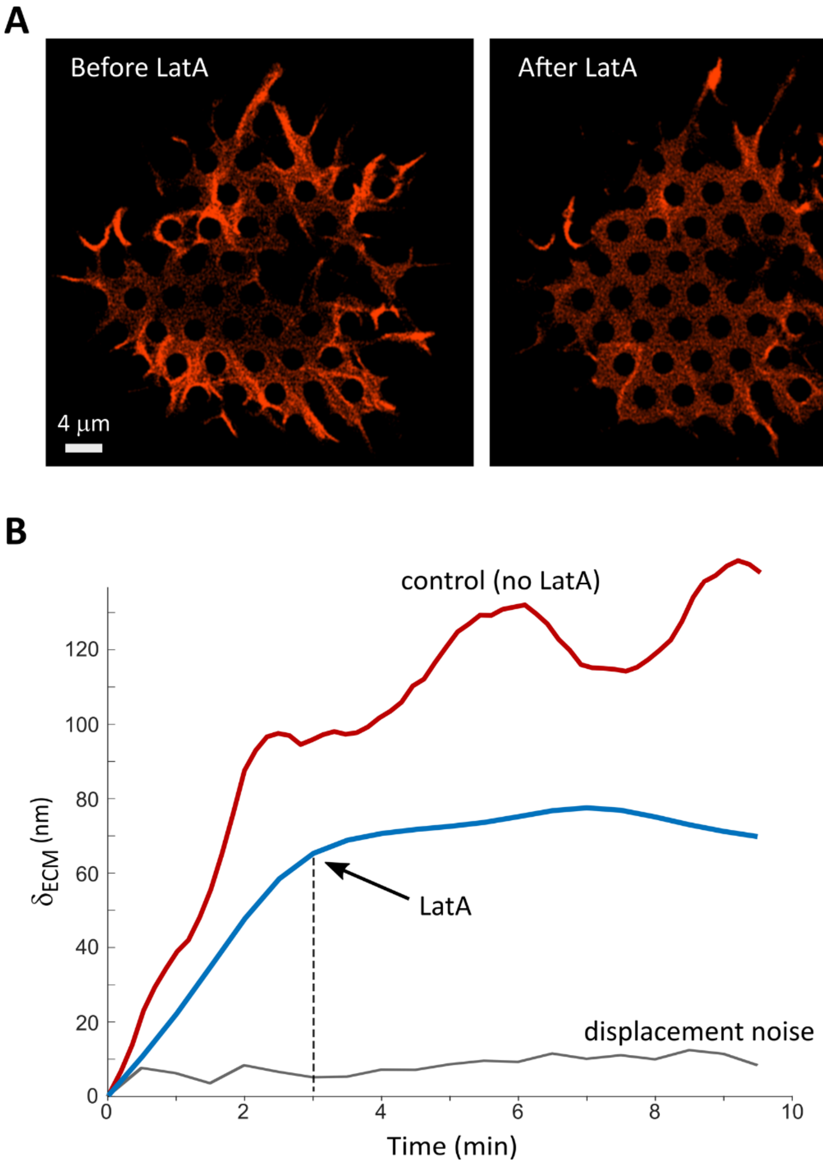


**Fig. S5.** Latrunculin A treatment blocks pillar displacement. (A) Images from a video of a MEF spreading on 2µm diameter pillars (stiffness = 31pN/nm) immediately before and immediately after addition of 330nM Latranculin A (LatA). Time difference between the two images is 30 s. (B) Example of a displacement curve of a single pillar (blue) whose movement was immediately blocked by LatA addition. Grey curve is the displacement noise (pillar that was not in contact with the cell throughout the experiment). Red curve is an example of pillar displacement by a cell that was not treated with LatA.

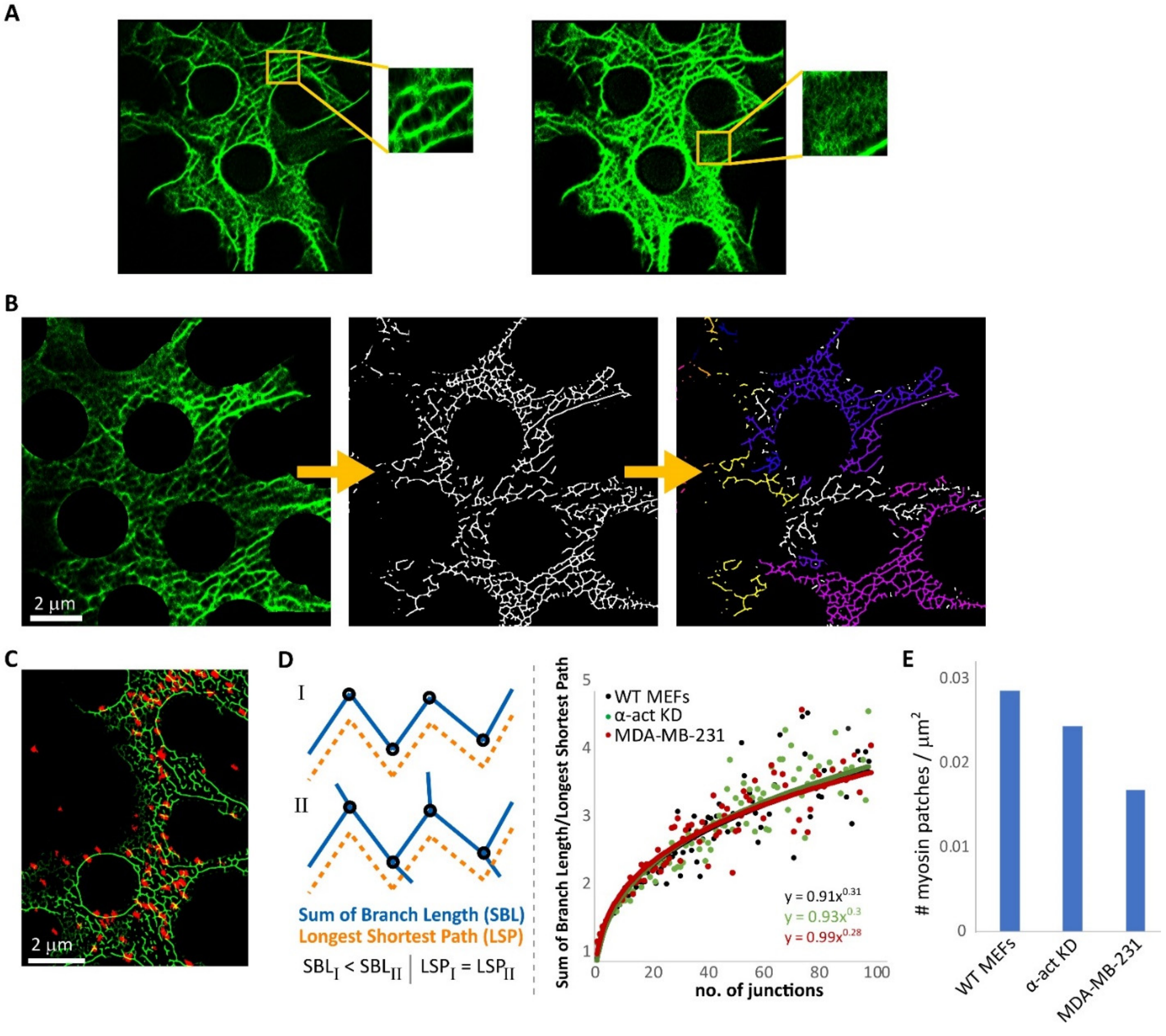


**Fig. S6.** Characterization of F-actin and p-myosin organization. (A) Images of a section of the cell edge on 2μm pillars stained for F-actin (bottom-left corner of the image is outside of the cell). The same image is presented twice, with decreased and enhanced brightness (left and right, respectively). The left zoom-in shows micron-scale actin structures whereas the right zoom-in shows nanometer-scale unstructured filaments. Box size in both cases is 1.7μm x 1.7μm. (B) Example of a super-resolution image of F-actin used to characterize the organization of the micron-scale actin structures between the pillars (top and right sides of the image are outside of the cell). Left: original image. Middle: segmented image. Right: analysis of the segmented image for continuous skeletons; each hue represents a single coherent skeleton. (C) Overlay of p-myosin (red) and F-actin (green) at the cell edge (right side of the image is outside of the cell). (D) Analysis of skeletons' complexity. We defined complexity as the ratio between the sum of branch length and the longest shortest path of each skeleton. In the example given on the left, both skeletons have the same shortest longest path; however, skeleton II, which is more complex, has a higher sum of branch length. Right: ratio of sum of branch length and the longest shortest path as a function of the number of junctions. No difference is observed between the three cell lines. For clarity purposes, each data point presented here is the average ratio for each no. of junctions (n>1500 skeleton all together, from >33 cells in each case, out of which >300 skeletons had 10 or more junctions). (E) Mean number of myosin clusters per μm$^2$ for each cell line.

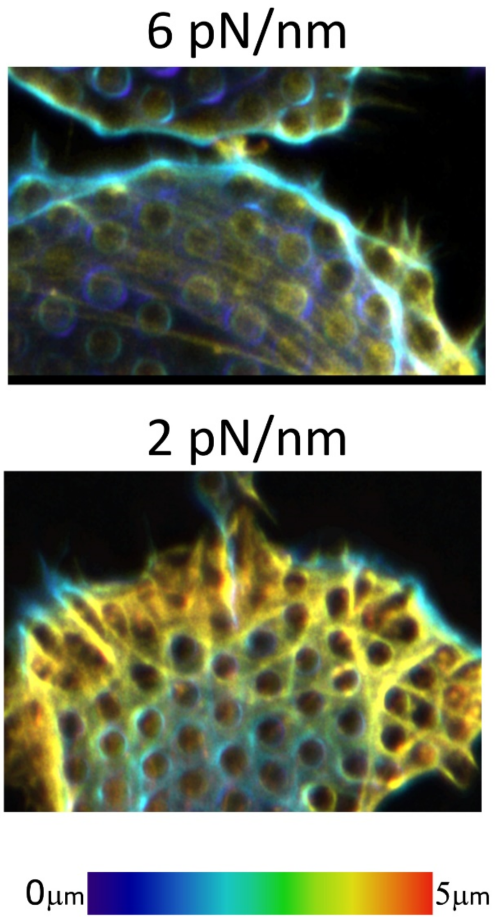


**Fig. S7.** Z-stack projections of the edges of REF52 cells on 2 and 6pN/nm pillars stained for F-actin color-coded for depth after 5h of spreading. F-actin around the pillars at the edge (blue hues) are connected to stress fibers 2-4μm above (yellow hues). Thick actin stress fibers (yellow-orange hues) are observed 2-4μm above the fibers directly surrounding the pillar (blue hues).

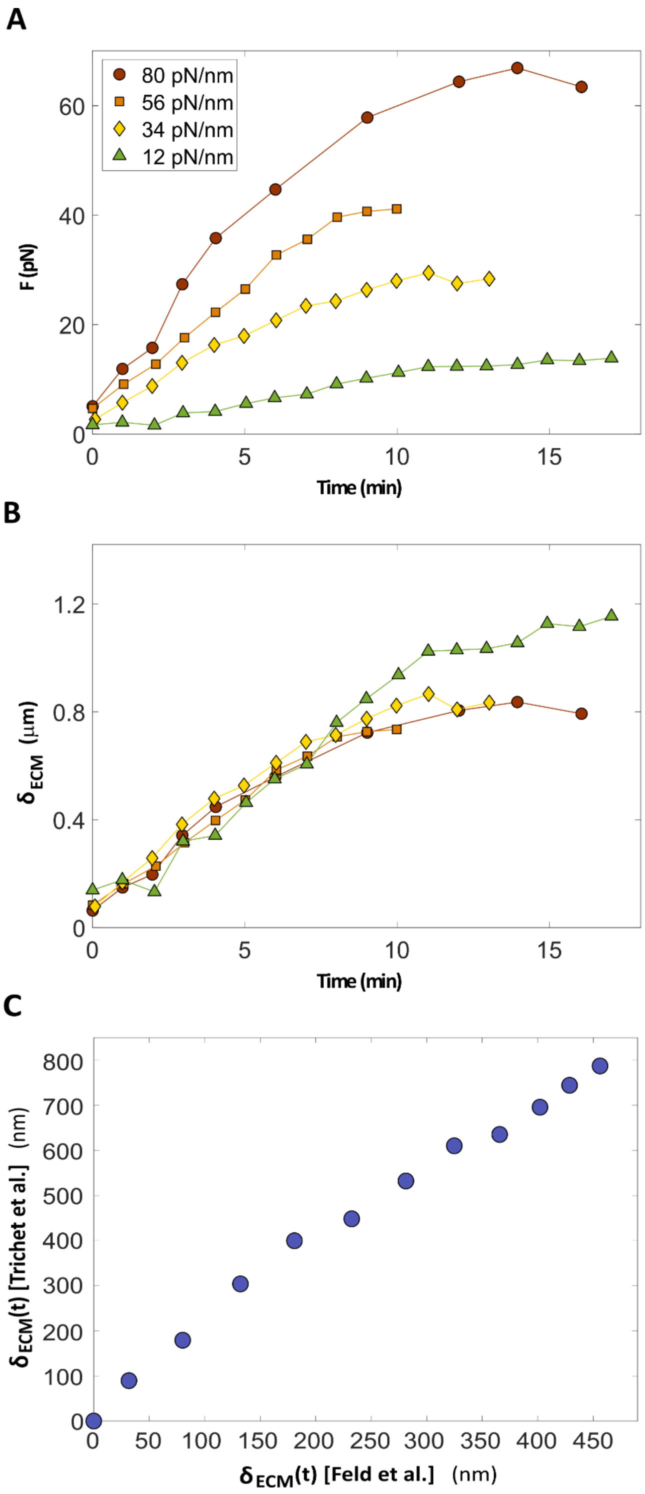


**Fig. S8.** (A) $F(t, k_{ECM})$ curves with different $k_{ECM}$ values were extracted from Fig. 2A of Trichet et al. Note that in the original publication, data for $k_{ECM} = 4.7$ pN/nm are also presented; however, the size of the symbols there is comparable to the actual measurement values, preventing reliable extraction of these data. (B) Replotting of the data in panel A in the rescaled form, $F(t, k_{ECM})/k_{ECM}$. (C) $\delta_{ECM}(t) = F(t, k_{ECM})/k_{ECM}$ of REF52 cells under well-spread steady-state conditions as measured in this work (Fig. 5B) vs. that measured by Trichet et al. (panel B here).

#### Supplementary Note 1: Relation to the two-spring model

The basic arrangement of elements in our model, which is sketched in Fig. 1B, is similar to the so-called two-spring model of Schwarz and co-workers (*1*) (cf. Fig. 4 and Sect. 4 therein). There, like in our model, myosin II motors displace the intracellular structures that are represented by a linear spring (with rigidity $k_{Act}$, in our notation), which are themselves connected in series to the extracellular matrix that is also represented by a linear spring (with rigidity $k_{ECM}$, in our notation). Yet, the two models are qualitatively and fundamentally different. In the two-spring model (*1*), the time rate of change of the contractile displacement $\dot{\Delta}(t)$ is assumed to follow a linearized Hill-like relation of the form $\dot{\Delta}(t) = v_0 \left(1 - {F(t)}/{F_s}\right)$, where the "free velocity" $v_0$ and the stall force $F_s$ are intrinsic cellular properties. As $F(t)$ depends on the rigidity $k_{ECM}$, so does $\dot{\Delta}(t)$ (which should in fact be written as $\dot{\Delta}(t, k_{ECM})$), in sharp disagreement with our findings. Consequently, it is not surprising that the predictions of the two-spring model (*1*) are qualitatively inconsistent with our results. In particular, taking the time derivative of Eq. (1) in the main text and using the expression for $\dot{\Delta}(t)$ to solve for $F(t)$, one obtains $F(t) = F_s \left(1 - e^{-t/t_K}\right)$ with $t_K = \frac{F_s}{v_0\, k_{ECM}}$ (cf. Eq. (9) in Schwarz et al. (*1*)). This prediction is qualitatively inconsistent with our findings, e.g. the timescale $t_K$ depends on the rigidity $k_{ECM}$.

#### Supplementary Note 2: Additional theoretical considerations and relations to existing literature

In our model, the actin structures are characterized by an effective rigidity $k_{Act}$. It is important to note that in general, rigidity (i.e. a spring constant) is not an intrinsic material property, but rather a combination of an intrinsic property (i.e. an elastic modulus) and a geometric property of the object of interest. This means that $k_{Act}$ also depends on the geometric properties (e.g. the typical thickness, length, and connectivity) of the actin structures, and since these are time-dependent, so is $k_{Act}$. Moreover, as the geometric properties of actin structures are known to be affected by the external rigidity (*2*), we expect $k_{Act}$ to depend on it as well. Taken together, one should in fact have $k_{Act}(t, k_{ECM})$, though in the manuscript we use just $k_{Act}$. Our assumption is that $k_{Act} \gg k_{ECM}$ at any point in time (otherwise $\delta_{ECM}$ cannot be higher than 0).

As explained in the main text, our model and its supporting evidence are consistent with other available observations in the literature, and hence provide a unifying picture of cellular contractility. Here we provide additional details about the relations between our work and existing literature. The prediction $F(t, k_{ECM}) \approx k_{ECM}\, \Delta(t)$ (Eq. (1) in the manuscript, once the $k_{ECM}$-independence of $\Delta(t)$ is taken into account), valid for $k_{Act}(t, k_{ECM}) \gg k_{ECM}$, is also consistent with the data of Saez et al. (*3*), where it has been shown that the contractile force at saturation, $F(t \to \infty, k_{ECM})$, is proportional to $k_{ECM}$, in agreement with our predictions (recall that $\Delta(t \to \infty)$ is finite). We would like to stress that while the proportionality between $F(t \to \infty, k_{ECM})$ and $k_{ECM}$ has been noted in both Saez et al. (*3*) and Trichet et al. (*4*) (in fact, in the latter it has been also explicitly noted that $dF(t = 0, k_{ECM})/dt$ is proportional to $k_{ECM}$), none of these works – and to the best of our knowledge no other work – has proposed that the contractile force $F(t, k_{ECM})$ is proportional to $k_{ECM}$ at any time $t$, which implies the existence of an intrinsic (non-mechanosensitive) time-dependent contractile displacement $\Delta(t)$.

In Ghibaudo et al. (*5*) the saturation force $F(t \to \infty, k_{ECM})$ has been measured over a wide range of rigidities $k_{ECM}$, also probing the high rigidity regime, $k_{Act}(t, k_{ECM}) \ll k_{ECM}$. In this regime, the force takes the form $F(t \to \infty, k_{ECM}) = \Delta(t \to \infty,\ k_{ECM})\, k_{Act}(t \to \infty,\ k_{ECM})$. As discussed extensively in the manuscript, $\Delta(t \to \infty,\ k_{ECM})$ is independent of $k_{ECM}$. Moreover, both actin structures (e.g. stress fibers) and focal adhesions are known to become independent of the ECM rigidity for large rigidities (*6*), hence we expect $k_{Act}(t \to \infty,\ k_{ECM})$ to be also independent of $k_{ECM}$, implying that the saturation force $F(t \to \infty, k_{ECM})$ is independent of $k_{ECM}$ for $k_{Act}(t, k_{ECM}) \ll k_{ECM}$. Consequently, we expect $F(t \to \infty, k_{ECM})$ to be proportional to $k_{ECM}$ for $k_{Act}(t, k_{ECM}) \gg k_{ECM}$ and to be independent of it for $k_{Act}(t, k_{ECM}) \ll k_{ECM}$, exactly as observed in Fig. 2 of Ghibaudo et al.(*5*).

Finally, note that in the $k_{Act}(t, k_{ECM}) \ll k_{ECM}$ regime, e.g. in experiments on cells adhering to glass plates, our model predicts $\delta_{Act}(t) \approx \Delta(t)$. In this regime, the active displacement generated by the myosin motors is accommodated by the actin structures (since the ECM cannot be deformed), resulting in *stretched* actin filaments, $\delta_{Act}(t) > 0$. This property leads to a clear prediction in relation to laser cutting/ablation experiments in which stress fibers of contractile adherent cells are cut with laser and their subsequent dynamics are tracked. In particular, it implies that if cells adhering to glass (i.e. essentially infinitely rigid) plates are first allowed to reach the well-spread steady state for which $\delta_{Act}(t \to \infty) \approx \Delta(t \to \infty) > 0$, upon cutting/ablation, stress fibers are expected to *instantaneously retract* by an amount $\Delta(t \to \infty)$. This prediction is fully consistent with the laser cutting/ablation experiments of Russell et al. (*7*), where cells have been allowed first to reach their well-spread steady-state while adhering to glass substrates. Upon laser cutting/ablation, stress fibers are observed to instantaneously retract by ~2 μm (see Fig. 1D in Russell et al. (*7*)), in agreement with the ~1μm pillar deflection observed in Supplementary Fig. 8B, which provides an estimate of $\Delta(t \to \infty)$ (a factor of 2 emerges from the fact that the stress fiber retraction effectively corresponds to releasing the deflection of 2 pillars, one from each edge). Overall, our model and the picture of cellular contractility emerging from it are consistent with a significant range of observations, corresponding to various experimental protocols.
